# Inhibitory inputs from hippocampal CA1 to retrosplenial agranular cortex gate social behavior

**DOI:** 10.1101/2022.08.09.503424

**Authors:** Yuhan Shi, Jingjing Yan, Xiaohong Xu, Zilong Qiu

## Abstract

Retrosplenial cortex has been implicated in processing sensory information and spatial learning, with abnormal neural activity observed in association with psychedelics and in mouse and non-human primate models of autism spectrum disorders (ASD). The direct role of the retrosplenial cortex in regulating social behaviors remains unclear. This work reveals that the neural activity of retrosplenial agranular cortex (RSA), a subregion of retrosplenial cortex, is initially activated, then quickly suppressed upon social contact. The up-down phase of RSA neurons is crucial for normal social behaviors. PV-positive GABAergic neurons in the hippocampal CA1 region were found to send inhibitory projections to RSA. Blocking these CA1-RSA inhibitory inputs significantly impaired social behavior. Notably, enhancing the CA1-RSA inhibitory input could rescue social behavior defects in an ASD mouse model. This work suggests a neural mechanism for salience processing of social behavior and identifies a potential target for ASD intervention using neural modulation approaches.

## INTRODUCTION

Social behaviors are essential for the survival and reproduction of mammals (Chen and Hong, 2018). The execution of social behaviors requires the perception of sensory information, salience processing of social-related information, and further integration in the prefrontal cortex (Kingsbury and Hong, 2020). Abnormal social behaviors linked to neuropsychiatric disorders, such as autism spectrum disorder(ASD), seriously affect individuals’ quality of life (Bourgeron, 2015; Iakoucheva et al., 2019; Vorstman et al., 2017).

Recently findings show that ketamine treatment increased neural activity in retrosplenial cortex (RSC) and decreased social behaviors in mice (Vesuna et al., 2020). Abnormal upregulation of neural activity was observed in the RSC of *MECP2*-overexpressing mouse and increased functional connectivity of RSC and other brain regions in the non-human primate models for ASD, compared to wild-type (WT) animals (Cai et al., 2020; Yu et al., 2020). Our group, alongside with Li and colleagues, found significant alterations in excitatory and inhibitory synaptic transmission in RSC in various genetic mouse models of ASD (Shang et al., 2021; Yang et al., 2021). These findings collectively suggest that the RSC plays a role in regulating social behavior.

The RSC consists of two subregions, the retrosplenial granular cortex (RSG) and retrosplenial agranular cortex (RSA), which are anatomically and functionally distinct (Aggleton et al., 2021; Sigwald et al., 2019; Vogt, 2009). The RSC receives synaptic inputs from several areas, including visual cortex, hippocampus, thalamic nuclei, and the anteroventral nucleus (Brennan et al., 2021; Murakami et al., 2015; Powell et al., 2020). In particular, synaptic inputs to the RSC from the hippocampus are crucial for memory formation and consolidation (Nitzan et al., 2020; Opalka and Wang, 2020; Yamawaki et al., 2019). Intriguingly, our findings indicate that neural activity of hippocampal CA1 neurons is causally correlated to social behaviors in MECP2 overexpression mice, an mouse model for ASD (Sun et al., 2020).

Salience processing of social versus non-social inputs from sensory systems is critical for interactive behaviors with conspecific partners during social-related inter-brain neural activity in the prefrontal cortex (Chen and Hong, 2018; Kingsbury and Hong, 2020). Consequently, we hypothesized that the neural circuit from the hippocampus to RSC plays a pivotal role in modulating social behaviors through salience processing of sensory inputs, which may be disrupted by ASD-related genetic mutations.

In this work, we discovered that neural activity of RSA neurons is initially activated but rapidly suppressed upon social contact. Interestingly, the up-down phase of RSA neurons is essential for social behaviors. We identified that PV-positive interneurons in the hippocampal CA1 region send long-range inhibitory projections to the RSA. Blocking these CA1-RSA inhibitory inputs significantly impaired social behavior. Remarkably, enhancing the CA1-RSA inhibitory input can rescue social interaction defects in an ASD mouse model. These findings suggest that the inhibition input from CA1 to RSA upon social contact serves as a salience control mechanism for social behavior progression, potentially by filtering out non-social sensory information in RSA regions. This work not only proposes a neural mechanism for salience processing of social information but also highlighsts a candidate brain region for ASD intervention using neural modulation approaches.

## Results

### Up-down phase of RSA neurons activity observed phase during social interaction

Initially, we examined whether RSC neurons may be activated during social interactions. Immunostaining for c-fos, an immediately early genes indicative of neural activity, was performed on brain slices from mice that interacted with either novel intruder mice or with objects for 1.5 hours in a home-cage experiment (Figure 1A-C). Mice exposed to novel intruders displayed a significantly increased c-fos level in RSA neurons compared to those exposed to objects, suggesting that RSA neurons are activated upon social interaction (Figure 1D-F).

**Figure 1.**
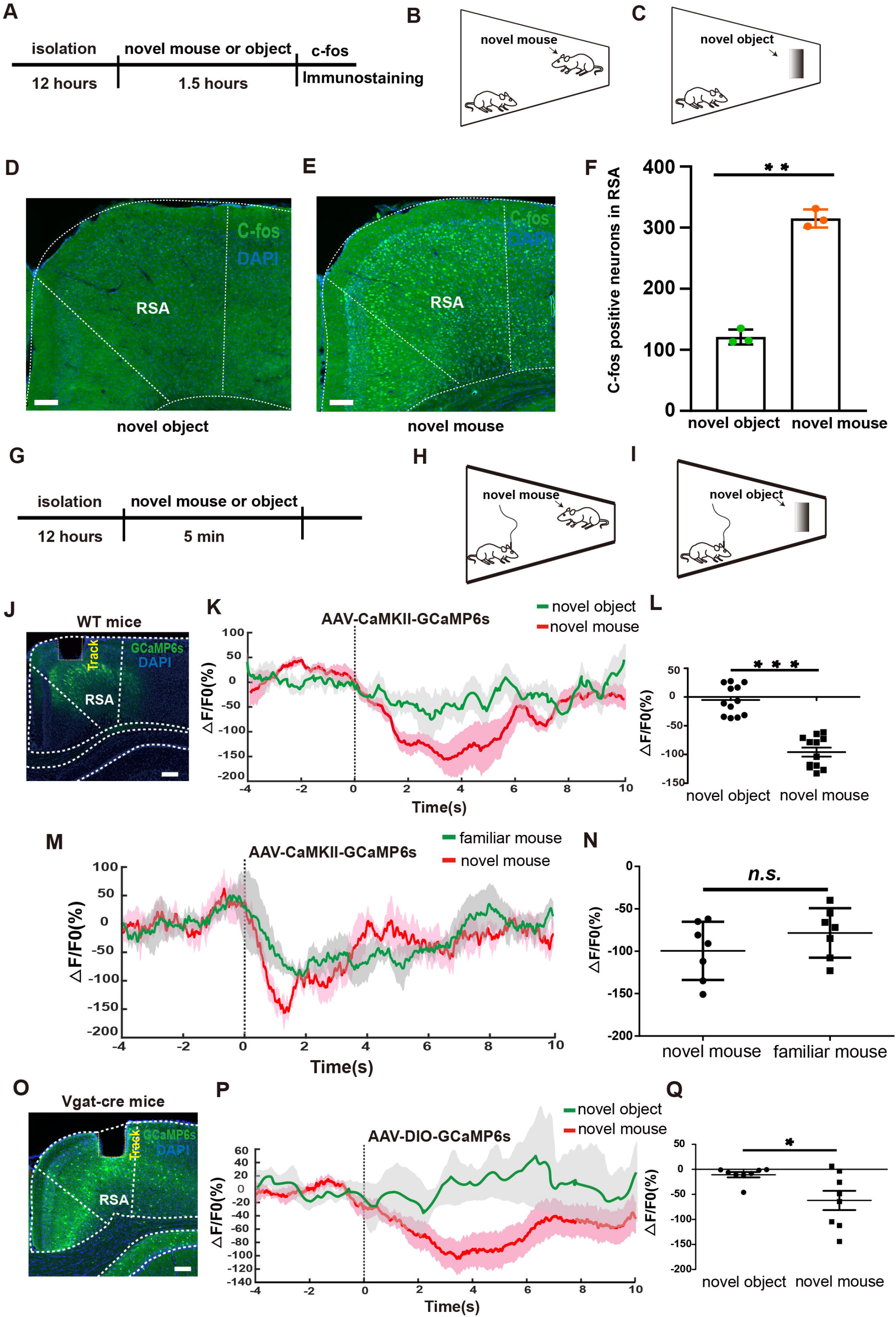
RSA is activated by social contact and inhibited after sniffing initiation. (A-C) Schematic illustration of the home-cage test. Representative images showing c-fos expression in RSA after interaction with object (D) and novel mice (E). Scale bar, 300μm. (F) Quantification of numbers of c-fos positive neurons in RSA from (D, E). (G-H) Schematic illustration of the home-cage test for fiber-photometry experiment. (J) Implantation of the optic fiber for fiber photometry in RSA layer IV-V of a WT mouse injected with AAV-CaMKII-GCaMP6s. Scale bar, 250μm. (K) Mean calcium transient associated with social interaction. Solid lines indicate mean and shadow areas indicate SEM (green: object, red: novel). Dash line at the 0s time point represents the time point when mice actively touched the novel mice with nose. (L) Quantification of average ΔF/F value of (0-6s) from mice with either objects or novel mice (baseline is the mean value of -4s-0s) (n = 12 mice for each group). (M) Mean calcium transient associated with social interaction process. Solid lines indicate mean and shadow areas indicate SEM (green: familiar mouse, red: novel mouse). Dash line at the 0s time point represents the time point when the mice actively touched the novel mice with nose. (N) Quantification of average ΔF/F value of (0-6s) from mice with either familiar or novel mice (baseline is the mean value of -4s-0s) (n = 7 mice for each group). (O) Implantation of the optic fiber for fiber photometry in RSA of the Vgat-cre mouse injected with AAV-DIO-GCaMP6s. Scale bar, 250μm. (P) Mean calcium transient associated with social interaction process. Solid lines indicate mean and shadow areas indicate SEM (green: object, red: novel). Dash line at the 0s time point represents the time point when the mice actively touched the novel mice with nose. (Q) Quantification of average ΔF/F value of (0-6s) from mice with either objects or novel mice (baseline is the mean value of -4s-0s) (n = 8 mice for each group). \**p* < 0.05, ** *p* < 0.01, *** *p* < 0.001. Error bars represent mean ± SEM.

The RSC is known to be involved in integrating sensory information(Alexander and Nitz, 2015; Vann et al., 2009; Wolbers and Buchel, 2005). To identify which RSC subregion may receive input from sensory cortices, we injected retrograde retroAAV-Cre-mCherry virus into RSA or RSG regions of Ai-9 (Rosa26-CAG-loxp-stop-loxp-tdTomato) mice (Figure S1A-D). Our findings revealed that RSA, rather than RSG, primarily received inputs from the primary visual cortex (V1), suggesting an essential role for RSA in relaying information to higher centers during social interaction. We further performed anterograde tracing by injecting AAV-hSyn-ChR2-mCherry into V1 (Figures S1E). A considerable number of ChR2-mCherry-expressing axon terminals were found in RSA, but not in RSG (Figures S1F), consistent with previous findings in rat (Aggleton et al., 2021).

Since c-fos protein levels persist for hours after being rapidly induced by neural activity, we aimed to investigate RSA neuron activity dynamics during social interactions. We injected AAV-CaMKII-GCaMP6s virus into RSA and implanted an optic recording fiber. With this setup, we recorded real-time Ca^2+^ dynamics in RSA neurons using fiber photometry in layer IV-V of RSA in WT mice while mice during home-cage interactions (Figure 1J). Interestingly, the calcium signal decreased immediately following a novel mouse sniff, compared to sniffing a novel object (Figure 1K, L). Moreover, Ca^2+^ signals in RSA rapidly declined after social contact, regardless of whether the mouse encountered a familiar or novel conspecific (Figure 1M, N), indicating that the decrease in neural activity in RSA is specific to social interaction in mice.

We then investigated whether local GABAergic interneurons might inhibit RSA neurons. To examine this possibility, we injected AAV-DIO-GCaMP6s virus into VGAT-ires-Cre mice, specifically labelling GABAergic neurons in RSA (Figure 1O). Notably, we observed a significantly decline in calcium transients in RSA GABAergic neurons within seconds after social contact (Figure. 1P, Q). Our results demonstrated that both excitatory and inhibitory neurons in RSA is quickly surpressed by a strong inhibitory input during sniffing, despite RSA neurons being initially activated upon social contact.

### Inhibition of RSA neurons after social contact promotes social behavior

To further investigate whether the up-down phase of RSA neurons is essential for social interactive behaviors, we employed optogenetics with the fine temporal resolution to manipulate neural activity. We unilaterally injected AAV-expressing channelrhodopsin2 (ChR2)-EYFP or Guillardia theta anion channel rhodopsin (GtACR)-mCherry into RSA for precise temporal control of RSA neuron activity (Figure 2A). After 12 hours isolation, we examined how manipulating neural activity in RSA affected social interactions between a home-cage and a novel mouse. When we constantly activated excitatory RSA neurons during social interaction by stimulating ChR2, mice displayed significantly decreased social interaction, as measured by cumulative sniffing time, interaction time, and interaction frequency, compared to AAV-EYFP-expressing mice (Figure 2B-D).

**Figure 2.**
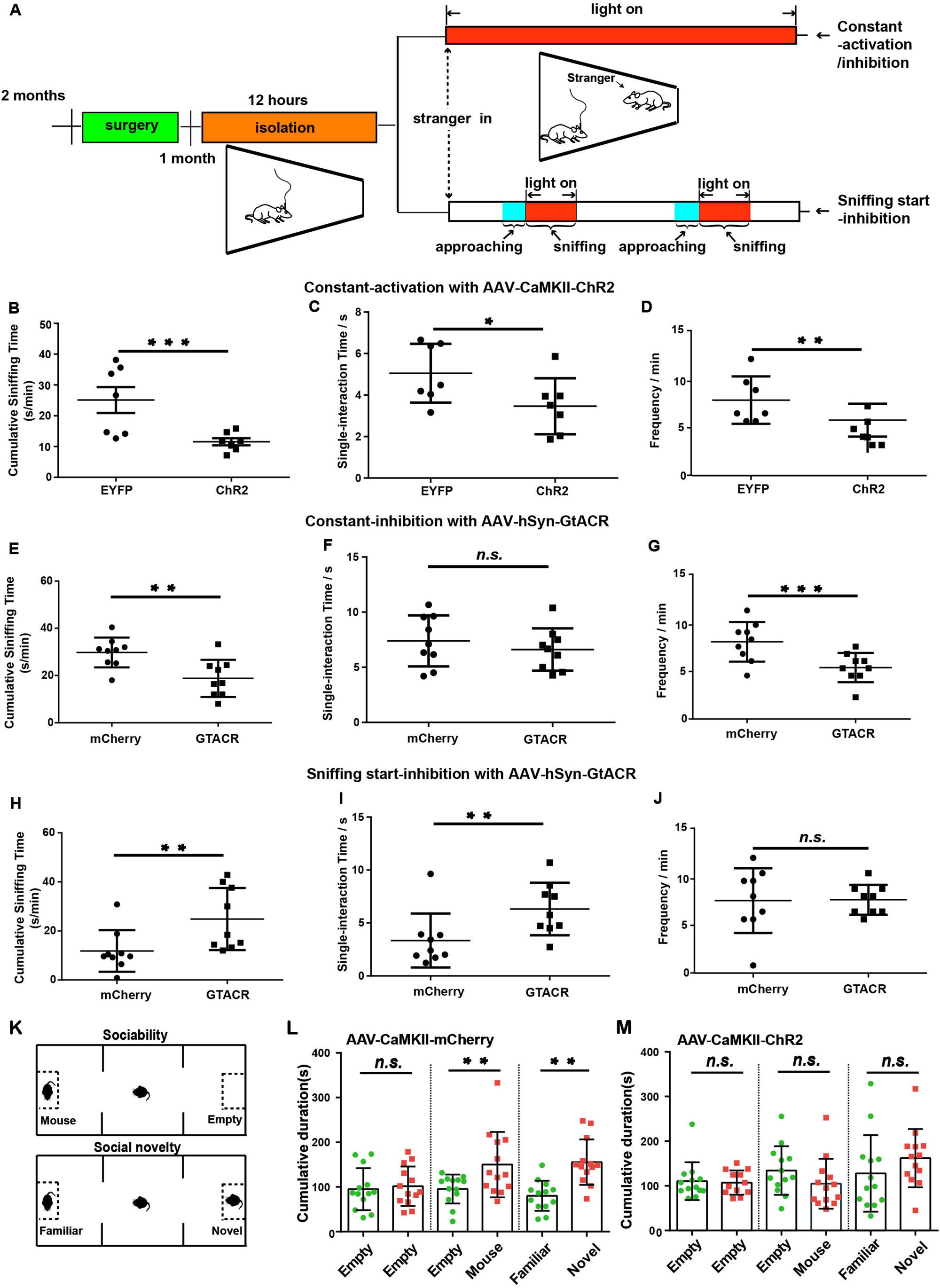
Sniffing start-inhibition of RSA facilitates the home-cage social behavior. (A) Schematic illustration of the optogenetic manipulation in RSA of mice injected with AAV virus during social interaction in home-cage tests. Quantification of cumulative sniffing time (B), single-interaction time (C) and sniffing frequency (C) induced by constant 473 nm light activation in the ChR2 group and the EYFP group (n = 7 mice for each group). Quantification of cumulative sniffing time (E), single-interaction time (F) and sniffing frequency (G) induced by constant 550nm light inhibition in the GtACR group, comparing to the mCherry group (n = 9 mice for each group). Quantification of cumulative sniffing time (H), single-interaction time (I) and sniffing frequency (J) induced by sniffing-start 550nm light inhibition in the GtACR group, comparing to the mCherry group (n = 9 mice for each group). (K) Schematic illustration of the three-chamber test, including sociability and social novel tests. Quantification of cumulative duration during social approach and social novelty sessions in the three-chamber test for the mCherry group (L) or the ChR2 group (M) (n = 13 mice for each group). \**p* < 0.05, ** *p* < 0.01, *** *p* < 0.001. Error bars represent mean ± SEM. Optogenetic stimulation parameters (20Hz, pulse width 5ms)

We then explored whether inhibiting RSA neurons would affect mouse social behaviors. Following AAV-hSyn-GtACR-mCherry injection into the RSA region, we could optogenetically inhibit RSA neurons by photostimulation at various stages of social interaction (Figure 2A). First, we induced constant inhibition by applying 550 nm light to RSA neurons. Interestingly, this resulted in significantly reduced cumulative sniffing time and sniffing frequency of mice with novel intruder mice, indicating that constant blockade of RSA neural activity negatively impacts social behaviors (Figure 2E-G). Next, we mimicked the down phase by inhibiting neural activity immediately after sniffing initiation (sniffing start-inhibition) (Figure 2A). Remarkably, both accumulative and single interaction times with novel mice increased significantly following sniffing start-inhibition (Figure 2H-J), suggesting that suppressing RSA neural activity immediately after social contact enhances social interactive behaviors.

Additionally, we examined whether constant RSA neuron activation would affect mouse social behavior in the classic three-chamber test paradigm (Figure 2K). Consistent with our previous findings, RSA neuron activation led to a complete loss of sociability and social novelty preference (Figure 2L, M). We also observed that continuous RSA activation did not alter mouse anxiety levels in the open-field test (Figure S2A, B). This data demonstrated that constant RSA activation strongly inhibits social interaction behaviors in mice.

In conclusion, we determined that constant activation or inhibition RSA neurons during social interaction impairs mouse social behaviors, whereas suppressing RSA neurons immediately after social contact is necessary for proper social interactive behaviors. Abnormalities in sensory information processing have been identified in ASD patients and animal models (Wiggins et al., 2009), and correcting genetic defects in the sensory system has been shown to rescue behavioral defects in ASD mouse models. We hypothesize that the crucial suppression step in RSA may represent a filtering process that blocks non-social information within sensory inputs during social interaction.

### PV-positive neurons in hippocampal CA1 project to RSA

We next explored the origin of inhibitory inputs to RSA neurons during social interaction. First, we conducted retrograde tracing by injecting cholera toxin B subunit (CTB) into RSA and observed numerous labeled neurons in dorsal CA1 (Figure 3A). We then performed immunostaining to identify the subtype of labeled neurons. Intriguingly, we discovered that approximately 38% of the PV-positive neurons in CA1 were labeled by CTB (Figure 3A), suggesting that PV-positive neurons in CA1 may project to RSA. We further confirmed the synaptic connection between PV-positive neurons in CA1 and RSA by injecting retroAAV-hSyn-Cre-mCherry into RSA and AAV-DIO-GFP into CA1 (Figures S3A, B), revealing that 23% of all the GFP positive neurons in CA1 were GFP and PV double-positive (Figures S3C-E).

**Figure 3.**
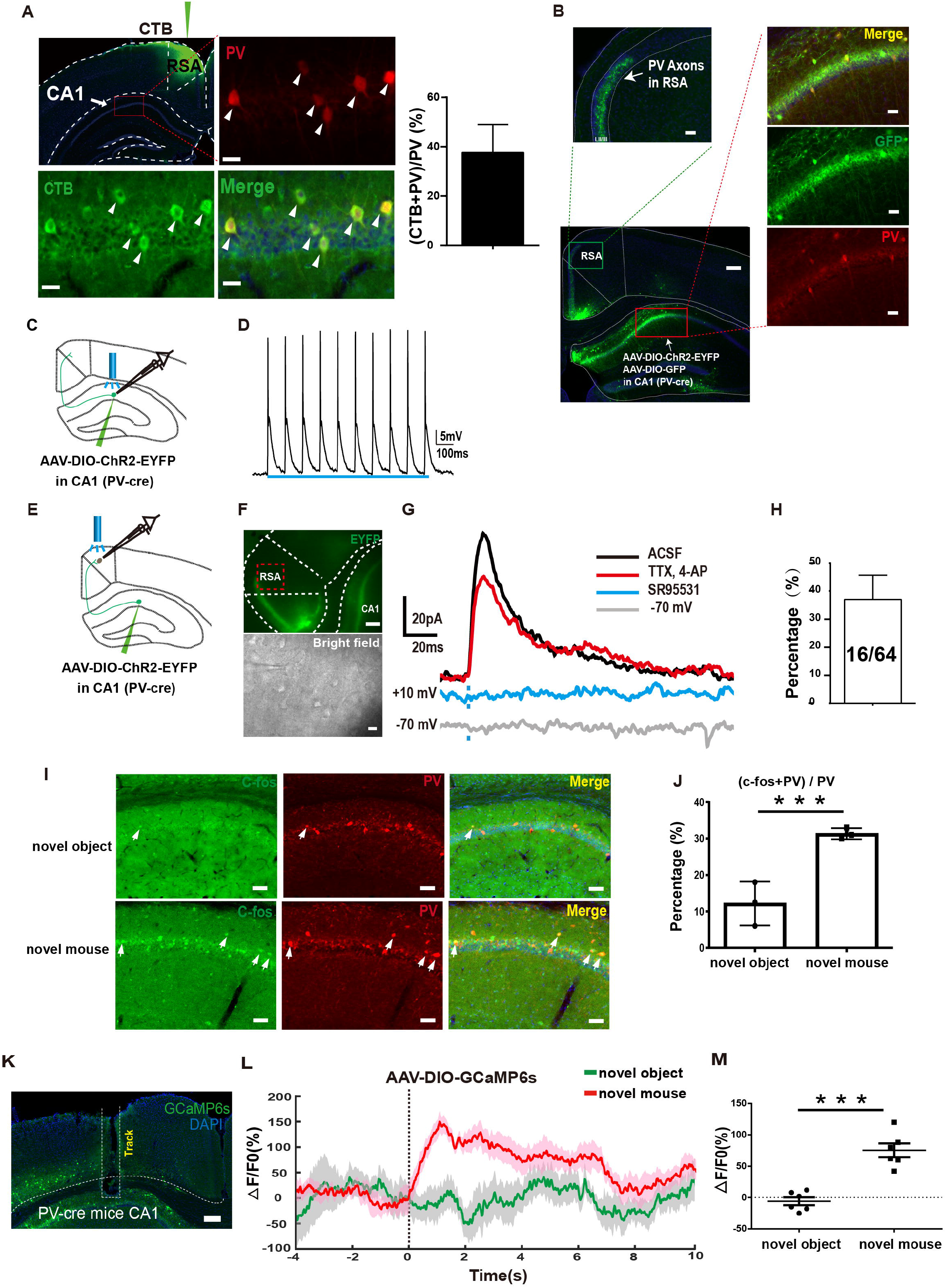
CA1 PV-positive neurons project to RSA and activated by social behavior. (A) Representative images showing retrograde tracing of RSA neurons with CTB. Quantification of the percentage of CTB and PV double positive neurons in total PV-positive neurons in CA1 (lower right panel). Scale bar, 750μm (upper left panel), 10μm (three zoomed-in areas of CA1). (B) Illustration of the axon terminal of the CA1 PV+ neurons in RSA (upper left panel), scale bar, 80μm; and the CA1 injection site of the of the virus; scale bar, 600μm (lower left panel), 20μm (three zoomed-in regions of CA1). (C) Schematic illustration of the patch-clamp recording on CA1 PV-positive neurons with activating the recorded neurons using 473 nm light in PV-cre mice injected with AAV-DIO-ChR2-EYFP into CA1. (D) The spikes recorded on EYFP-labeled neurons in CA1 of PV-cre mice with activated by stimulation of 473 nm light. (E) Schematic illustration of the patch-clamp recording on RSA neurons and simultaneously activating the CA1 PV axon terminal using 473 nm light. (F) Representative images of the patch-clamp recording under the infrared microscope illuminated by epifluorescence (upper panel), scale bar, 400μm and infrared-differential interference contrast illumination (bright field), scale bar, 10μm. (G) The representative trace (upper) of the recorded IPSCs on RSA neurons activated by 473 nm light with a +10mV holding potential which was totally abolished by GABA_A_ receptor antagonist SR95531 (gray trace), and partially blocked by TTX and 4-AP (red trace). The representative trace recorded on RSA neurons activated by 473 nm light with a -70mV holding potential (lower). (H) The evoked IPSCs were identified on 16 neurons of 64 neurons recorded from 5 mice. (I) Representative images of c-fos expression in CA1 PV+ neurons of mice with objects (upper three panels) and novel mice (lower three panels) in the home-cage test. Scale bar, 20μm. (J) Quantification of percentage of PV and c-fos double positive neurons in total PV-positive neurons in mice with new objects or with novel mice (n= 18 slices from 3 mice of each group). (K) Implantation of the optic fiber for fiber photometry in CA1 of PV-cre mouse injected with AAV-DIO-GCaMP6s. Scale bar, 250μm. (L) Mean calcium transient associated with social interaction process. Solid lines indicate mean and shadow areas indicate SEM (green: novel object, red: novel mouse). Dash line at the 0s time point represents the time point when the mice actively touched the novel mice with nose. (M) Quantification of average ΔF/F value of (0-6s) from mice with either novel object or novel mice (baseline is the mean value of -4s-0s) (n = 6 mice for each group). \*\*\**p* < 0.001. Error bars represent mean ± SEM.

To label axon terminals and soma, we injected Cre-dependent AAV-DIO-ChR2-EYFP and AAV-DIO-GFP into CA1 of PV-Cre mice (Figure 3B). ChR2-expressing axons were observed in layer II-III of RSC neurons, indicating that PV-positive neurons sent extensive synaptic termini to RSC (Figure 3B). We verified this observation by demonstrating that almost all of GFP-positive neuron colocalized with PV signals.

To examine whether CA1 PV-positive neurons form functional synaptic connections with RSA neurons, we utilized an optogenetic-electrophysiological approach to examine synaptic currents in brain slice preparations. After injecting Cre-dependent AAV-DIO-ChR2-GFP virus into the CA1 of PV-Cre mice, we performed whole-cell recording on layer IV-V neurons of RSA (Figure 3C). Wide-field photostimulation at 473 nm on ChR2-expressing axons revealed that RSA neurons exhibited outward inhibitory postsynaptic currents (IPSCs) at a command voltage of -10 mV, but no inward excitatory postsynaptic currents (EPSCs) when held at −70 mV (Figure 3D, E). Moreover, the IPSC could be fully blocked by the GABA_A_ receptor antagonist SR95531, indicating GABA_A_ receptor mediation (Figure 3E). We detected IPSCs in 16 out of 64 recorded neurons (5 mice, Figure 3F). We then performed whole-cell recording from ChR2-EYFP expressing neurons in CA1 with photostimulation (Figure 3G, Figure S3F) and observed the fast-spiking pattern characteristic of the PV-positive GABAergic neurons, confirming the specificity of genetic labeling in mice (Figure 3H).

We next assessed whether PV-positive neurons in CA1 were responsive to social interaction by performing immunostaining with c-fos in brain slices from mice interacting with either novel objects or novel mice. We found that the c-fos level of PV-positive neurons in CA1 was significantly higher than in mice with novel partners compared to those with novel objects (Figure 3I, J).

Additionally, we evaluated the responsiveness of CA1 PV-positive neurons to social interaction by injecting AAV-DIO-GCaMP6s virus into CA1 of PV-cre mice (Figure 3K). We observed that calcium signals in CA1 PV-positive neurons significantly increased when mice interacted with novel mice compared to novel objects, indicating that PV-positive neurons in CA1 specifically respond to social interaction (Figure 3L, M). Thus, we identified that a significant portion of PV-positive neurons in CA1 project to RSA and are activated by social stimulus. We hypothesize they may play a critical role in inhibiting the RSA neurons during social behavior.

The ventral CA1 has been implicated in social memory (Okuyama et al., 2016). In particular, PV-positive neurons in ventral CA1 have been found to play a pivotal role in social memory (Deng et al., 2019). Therefore, we sought to determine whether PV-positive neurons in ventral CA1 may also project to RSA. Initially, we performed the retrograde tracing by injecting CTB in RSA and found no labeled neuron in the ventral CA1 region (Fig. S3G, S3H). We then injected AAV-DIO-ChR2-EYFP into the ventral CA1 of PV-cre mice and observed no labeled terminals in the RSA and RSG region (Fig. S3I, S3J). We conclude that PV-positive neurons in ventral CA1 do not project to retrosplenial cortex(Deng et al., 2019; Okuyama et al., 2016).

### The inhibitory input from CA1 PV neurons to RSA is crucial for social behavior

To investigate the role of the inhibitory input from CA1 PV-positive neurons to RSA for social behavior, we employed a chemogenetic approach by injecting AAV-DIO-mCherry or AAV-DIO-hM4D into CA1 of PV-Cre mice and implanting a cannula into the superficial layer of RSA for local hM4D activation via clozapine-N-oxide (CNO) infusion (Figure 4A, B). Simutaneously, we injected AAV-CaMKII-GCaMP6s into RSA and implanted an optical fiber in layer IV-V of RSA to monito the neural activity in RSA (Figure 4B). We first tested the efficacy of hM4D-mediated inhibition through intraperitoneal CNO injection and observed the complete abolishment of the downward phase of calcium signals in RSA following social contact in the hM4D groups, compared to the mCherry group (Figure S4A, B).

**Figure 4.**
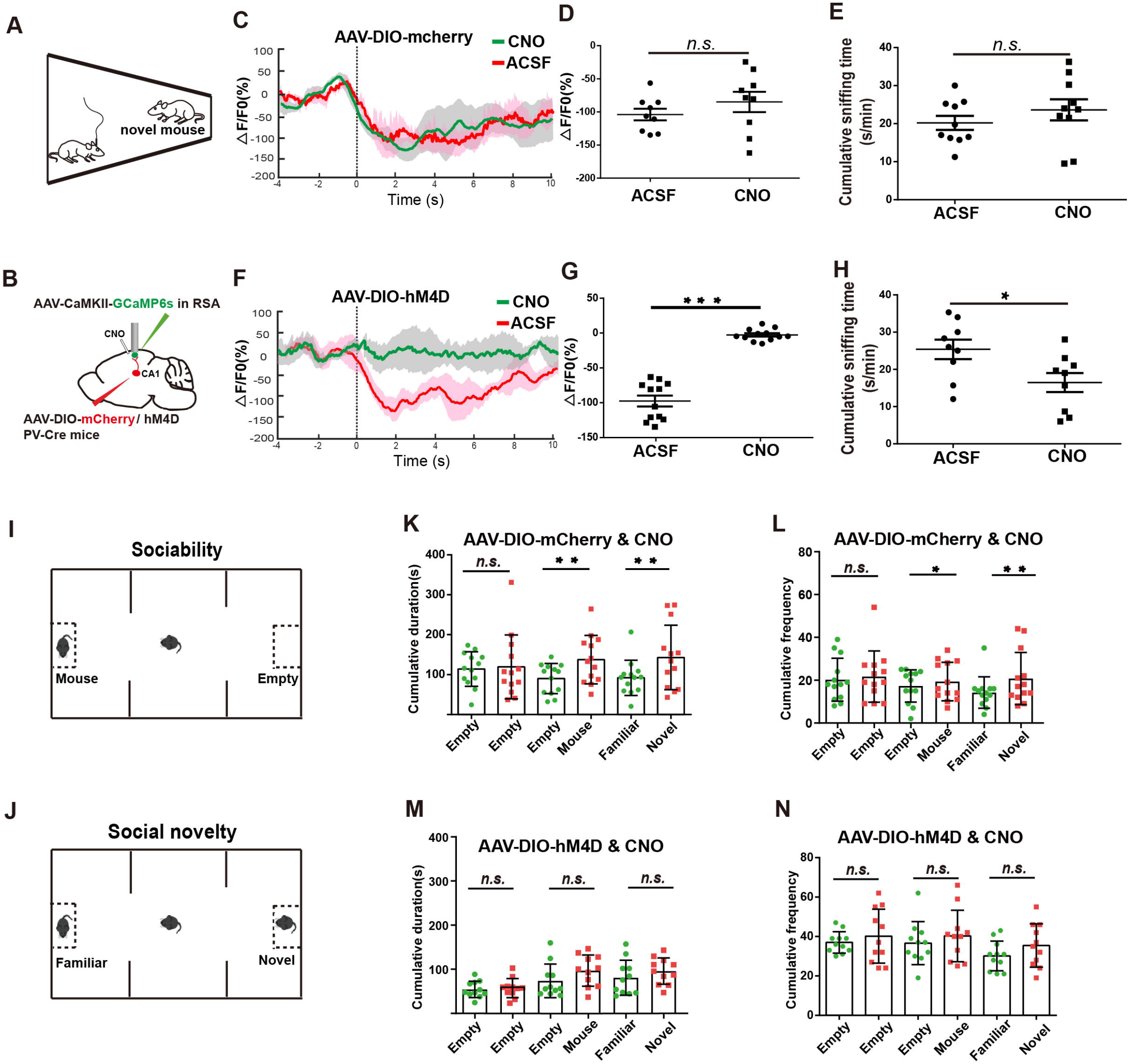
Blockade of the inhibitory projection from PV-positive neurons in CA1 to RSA abolished the inactivation in RSA during social behavior and led to social deficit. (A, B) Schematic illustration of set-up for the home-cage test and the pharmacogenetic manipulation with fiber-photometry recording on RSA excitatory neurons. AAV-DIO-mCherry/hM4D were injected into CA1 of PV-Cre mice and AAV-CaMKII-CCaMP6s were injected to RSA, with implantation of the optic recording fiber and pipette for CNO infusion. (C) Calcium transient of mice injected with AAV-DIO-mCherry during social interaction process (n= 9 mice for each group). Solid lines indicate mean values and shadow areas indicate SEM (green: CNO, red: ACSF). Dash line in the 0s time point represents the time point when the experimental mice actively touched the novel mice with nose. (D) Quantification of average ΔF/F value of 0-6s (baseline is the mean value of -4s-0s) from mice in either the ACSF group and the CNO group (n = 9 mice for each group). (E) Quantification of the cumulative sniffing time in the ACSF group and the CNO group in home-cage test (n = 10 mice for each group). (F) Calcium transient of mice injected with AAV-DIO-hM4D during social interaction process (n= 12 mice for each group). Solid lines indicate mean value and shadow areas indicate SEM (green: CNO, red: ACSF). (G) Quantification of average ΔF/F value of 0-6s (baseline is the mean value of -4s-0s) from the ACSF group and the CNO group (n = 9 mice for each group). (H) Quantification of the cumulative sniffing time in the ACSF group and the CNO group in home-cage tests (n = 9 mice for each group). Schematic illustration of sociability (I) and social novelty (J) session of the three-chamber test. Quantification of social time (K) and frequency (L) in the three-chamber test for in the mCherry group injected with CNO (n = 13 mice for each group). Quantification of social time (M) and frequency (N) in the three-chamber test for the hM4D group injected with CNO (n = 11 mice for each group). * *p* < 0.05, ** *p* < 0.01, *** *p* < 0.001. Bars represent mean ± SEM.

During the home-cage test, we locally injected CNO into RSA via the implanted cannula and recorded calcium transients of RSA neurons during social interaction (Figure 4A, B). Consistently with previous findings, the decline phase of calcium transients during social interaction was completely blocked in the hM4D-expressing group, but not in the mCherry-expressing group (Figure 4C, D, F, G). Surprisingly, social interaction was significantly impaired by blocking the inhibitory input from CA1 PV-positive neurons to RSA in the hM4D-expressing group, compared to the mCherry-expressing group (Figure 4E, H).

To further investigate whether the CA1-PV-RSA inhibitory projection influences social behaviors, we performed the classic three-chamber test to measure sociability and social novelty preference (Figure 4I, J). After CNO treatment, we measured the cumulative duration and frequency of social interaction and found that both sociability and social novelty preference were significantly impaired in the hM4D-expressing group, but not in the control group expressing only mCherry (Figure 4K-N).

These data indicate that inhibitory input from CA1 PV positive neurons to RSA contributes significantly to the suppression of RSA neurons following social contact and is essential for proper social interactive behaviors.

### Activation of the PV^+^ inhibitory projection from CA1 to RSA rescues the social deficit in an ASD mouse model

We next investigated whether enhancing the inhibitory projection from CA1 PV-positive neurons to RSA could rescue social deficits in ASD mouse models. Mutations of *MEF2C* have been identified in individuals with ASD and intellectual disabilities (Gilissen et al., 2014). *Mef2c* haploinsufficient (*Mef2c*^+/-^) mice exhibited ASD-like behaviors, such as defects in social interaction (Tu et al., 2017). We investigated whether enhancing the CA1 PV-RSA inhibitory pathway could rescue the social deficit phenotype of *Mef2c*^+/-^ mice. We first confirmed previous findings showing that *Mef2c*^+/-^ mice indeed exhibited social deficits in the home-cage experiment (Figure 5A-C). Additionally, the population of PV-positive neurons decreased in CA1 of *Mef2c*^+/-^ mice (Figure 5D, E), as reported (Tu et al., 2017).

**Figure 5.**
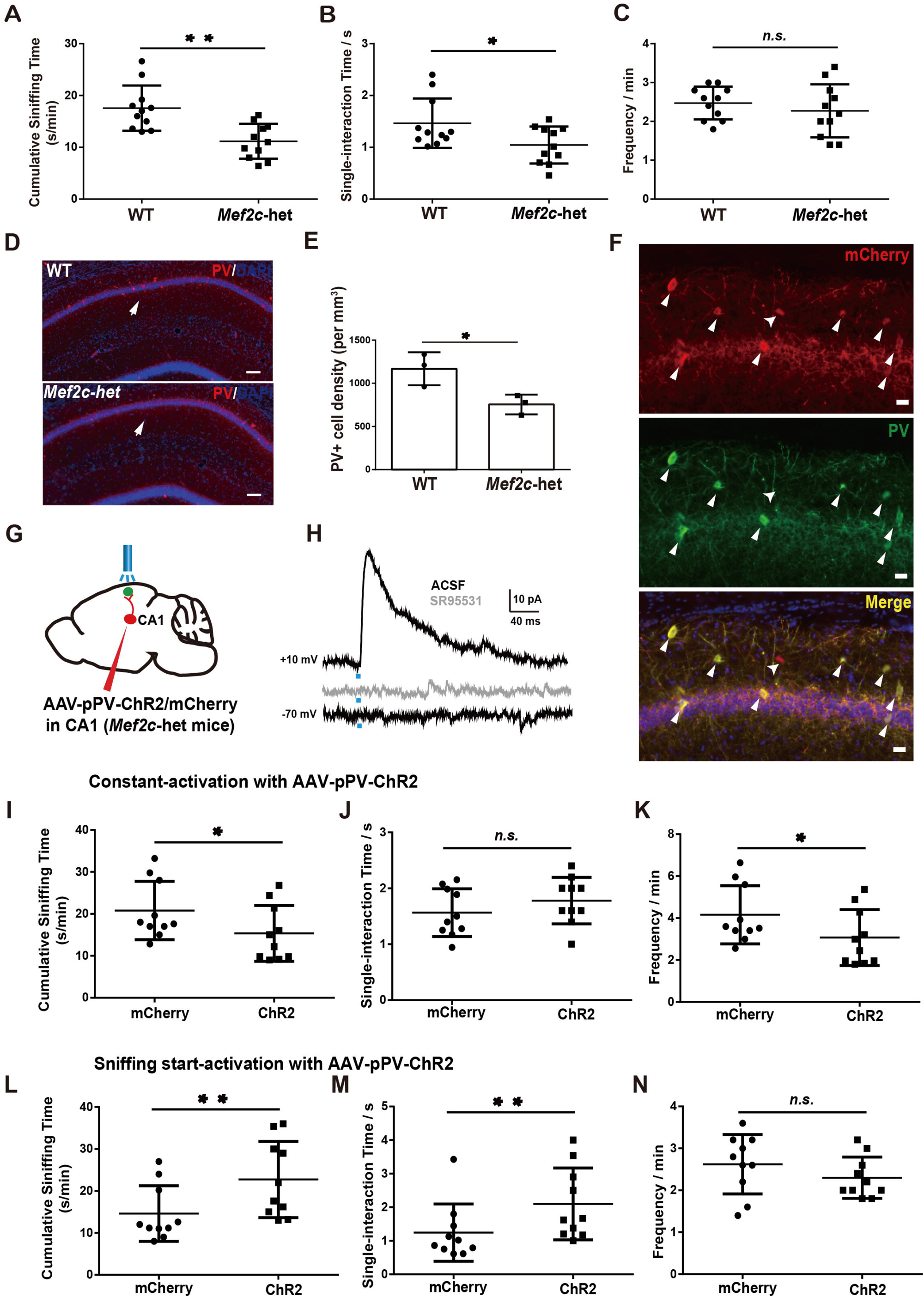
Activating the inhibitory projection from CA1 PV-positive neurons to RSA after sniffing initiation rescued the social deficit in *Mef2c*^+/-^ mice. Quantification of cumulative sniffing time (A), single-interaction time (B) and sniffing frequency (C) in the *Mef2c*^+/-^ mice group (*Mef2c*-het) and in the WT littermate group (WT). (D) Representative images showing decrease of the PV-positive neurons in *Mef2c*-het mice (lower panel); scale bar, 50μm. (E) Quantification of the PV-positive neurons in CA1 from the WT mice and the *Mef2c*-het mice. (F) Immunostaining of PV and mCherry in hippocampal slices of *Mef2c*-het mice injected with AAV-PV-ChR2-mCherry. Scale bar, 10μm. (G) Schematic illustration for optogenetic manipulation of PV-positive input to RSA in *Mef2c*-het mice. AAV-pPV-ChR2-mCherry was injected at CA1 of *Mef2c*-het mice with implantation of optic stimulation fiber in RSA. (H) The representative trace (upper) of the recorded IPSCs on RSA neurons activated by 473 nm light with a +10mV holding potential which was abolished by GABA_A_ receptor antagonist SR95531(gray trace), and the representative trace recorded on RSA neurons activated by 473 nm light with a -70mV holding potential (lower). Quantification of cumulative sniffing time (I), single-interaction time (J) and sniffing frequency (K) induced by constant 473 nm light activation in the *Mef2c*-het mice injected with either AAV-pPV-ChR2 (ChR2) or the AAV-pPV-mCherry (mCherry). Quantification of cumulative sniffing time (L), single-interaction time (M) and sniffing frequency (N) induced by sniffing-start 473 nm light activation in the *Mef2c*-het mice injected with either the AAV-pPV-ChR2 (ChR2) or the AAV-pPV-mCherry (mCherry). * *p* < 0.05, ** *p* < 0.01. Error bars represent mean ± SEM.

To specifically target PV-positive neurons in the hippocampal CA1 region of *Mef2c*^+/-^ mice, we used a PV-specific enhancer to drive expression of either ChR2 (AAV-pPV-ChR2) or mCherry (PV-mCherry) carried by AAV (Figure 5F) (Vormstein-Schneider et al., 2020). Through immunostaining with excitatory marker CaMKII and inhibitory marker parvalbumin, we found that the PV-specific enhancer faithfully labeled 85% of PV-positive neurons, and no excitatory neurons were labeled (Figure S5A-I).

To confirm whether ChR2-labeled PV-positive neurons in CA1 of *Mef2c*^+/-^ mice projected to RSA, we performed whole-cell recording of RSA neurons with 473 nm photostimulation (Figure 5G). We discovered that photo-activation ChR2-expressing axons evoked a robust IPSC in RSA neurons post-synaptically in brain slices, which could be completely blocked by SR95531, the antagonist of GABA_A_ receptor (Figure 5H). Among 20 recorded RSA neurons, we found 6 responsive neurons (Figure S6A).

We first examine whether manipulation CA1-PV-RSA circuits might affect social behaviors in WT mice. We injected AAV-pPV-ChR2 into CA1 of WT mice and photostimulated the axonal terminal in the RSA (Fig. S6B). We found that constant-activation of CA1-PV-RSA projection during social interaction led to decrease of sniffing time of WT mice with another mouse, in consistent with the previous finding (Fig. S6C-E). The sniffing start-activation of CA1-PV-RSA projection after social contact cannot further increase the social interaction time in WT mice (Fig. S6F-H), suggesting that the CA1-PV-RSA projection works properly in the WT mice during social interactive behaviors.

Since the number of PV-positive neurons in CA1 of *Mef2c*^+/-^ mice is significantly lower than that in WT mice (Figure 5D, E), we hypothesized that enhancement of CA1-PV-RSA projection with an optogenetic approach could rescue the social defects of *Mef2c*^+/-^ mice. Finally, we performed photostimulation in RSA of *Mef2c*^+/-^ mice during the home-cage test. We observed that overall duration and frequency of social interaction decreased further in the PV-ChR2 group of *Mef2c*^+/-^ mice during constant activation sessions, compared to the mCherry-expressing group (Figure 5I-K). Remarkably, social interactive time was significantly rescued in *Mef2c*^+/-^ mice if photostimulation to activate ChR2 was given right after sniffing started (Figure 5L-N). These results demonstrate that activation of the inhibitory projection of PV-positive neurons from CA1 to RSA during social contact is sufficient to rescue the impairment of social interaction in *Mef2c*^+/-^ mice.

## Discussion

Previously, our work and others demonstrated that synaptic transmission in retrosplenial cortex (RSC) neurons is disrupted in several ASD mouse models, including *Senp1*^+/-^ and *Fmr1*^-/y^ mice (Shang et al., 2021; Yang et al., 2021). Vesuna et al. showed that abnormally elevated neural activity in RSC of mice caused by ketamine treatment led to dissociated behaviors, including defects in social interaction in a dose-dependent manner (Vesuna et al., 2020). This evidence strongly suggests that the RSC plays a critical role in animal social interaction.

In this work, we proposed a model in which the inhibitory projection from PV-positive neurons of CA1 to RSA serves as a salience processing node to filter out non-social information flowing through RSA from sensory cortices. Although the processing of social information within RSA requires further investigation, the data presented here revealed neural mechanisms underlying the salience processing of social interaction behaviors. Since the hippocampus is one of brain regions extensively receiving sensory inputs besides sensory modules, we further hypothesized that the PV-positive neurons in the hippocampal CA1 region may serve as a salience processing switch by inhibiting non-social sensory information in RSA upon social interaction, even though all RSC neurons are activated upon social contact.

Social behavior is one of the fundamental interactive behaviors in mammals, which can be called “interactive behaviors within community” in non-human mammalian species. Within a community, the valence of social behavior is primarily influenced by social hierarchy and mating possibilities, while the salience of social behavior is largely determined by how the brain processes social vs. non-social information. The processing of social-related information salience is a crucial step for progression of social interactive behaviors. Furthermore, the salience of social interaction is clearly compromised in people with ASD, a group of brain disorders with strong genetic predispositions (Sandin et al., 2017).

Our findings suggest that defects in salience processing in RSA may be the causative factor for abnormal social behaviors in ASD patients. Clinically, defects in sensory perception, including visual and auditory systems, are widely observed in people with ASD. However, whether the defect in sensory perception is one of the causes or consequences of ASD remains debatable (Wiggins et al., 2009). This work suggests that defects in salience processing in RSA may be the causative factor for leading to autistic-like behaviors, as enhancing this pathway can rescue behavioral defects in mouse models for ASD.

The input and output of RSA circuits are quite intriguing. Although ventral CA1 was reported to play a critical role in social memory, we found that ventral CA1 did not project to RSA at all, suggesting PV-positive neurons in dorsal CA1 may be implicated in specific circuits governing social behaviors. Further work involves identifying the upstream of CA1-PV^+^ neurons, as well as the downstream of RSA neurons. The most critical question is to determine how social-related information is processed when the RSA is generally repressed during social interaction. More in-depth single-neuron resolution imaging may provide more insightful answers.

RSC in the human brain is easily accessible with neural modulation approaches, such as transcranial magnetic stimulation. Our finding opened an important venue through which one may test this hypothesis by manipulating neural activity in RSC in people with ASD and explore potential intervention methods.

## Availability of data and materials

The datasets used and/or analyzed in the current study are available from the lead contact on the reasonable request.

## Acknowledgements

We thank members of NPC for providing valuable comments for the manuscript. This work was supported by grants from the NSFC Grants (#81941015, #82021001); Strategic Priority Research Program of the Chinese Academy of Sciences (XDB32060202); Program of Shanghai Academic Research Leader, and the Science and Technology Commission of Shanghai Municipality (#2018SHZDZX05), Z.Q. is supported by GuangCi Professorship Program of Ruijin Hospital Shanghai Jiao Tong University School of Medicine.

