## Supplementary figures and methods for "Inhibitory inputs from hippocampal CA1 to retrosplenial agranular cortex gate social behavior"

Yuhan Shi, Jingjing Yan, Zhifang Chen, Xiaohong Xu, Zilong Qiu*

**Supplemental figures and legends, Figure S1-4**
**Supplemental methods and materials.**

#### Figure S1

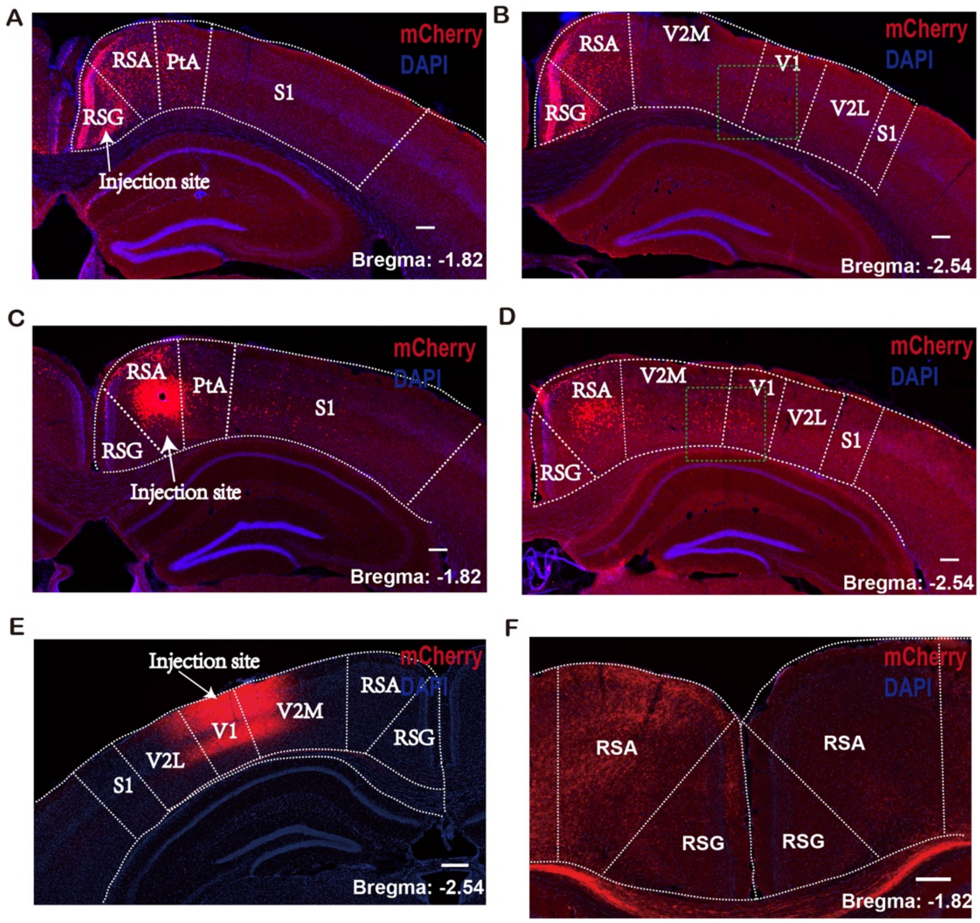


**Figure S1 RSA mainly receives input from primary visual cortex**
Representative images showing injection sites in RSG (A, B) and RSA (C, D) of retroAAV-cre-mCherry in Ai-9 mice; scale bar, 500µm. The mCherry-labeled neurons

are found in V1 of RSA-injected mice (D).

(E) Illustration of the injection site in visual cortex (V1) with AAV-hSyn-ChR2-mCherry; scale bar, 600µm.

(F) The ChR2-mCherry labeled axon terminals are mainly found in RSA; scale bar, 500µm.

### Figure S2

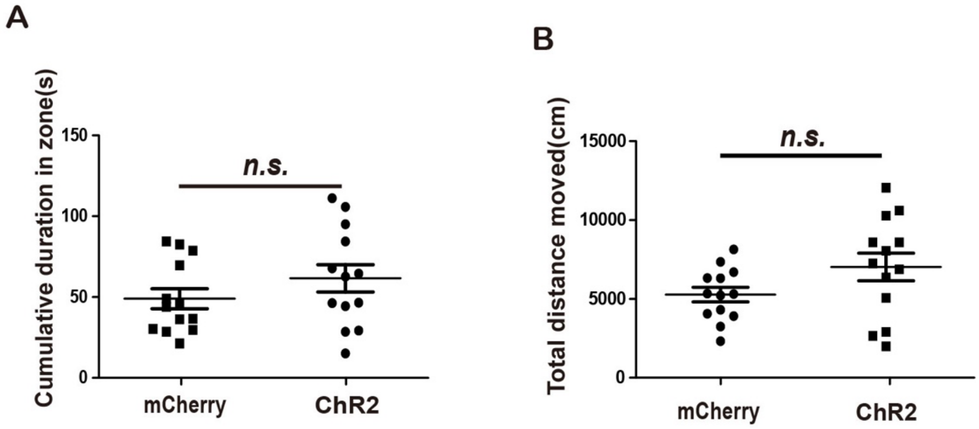


**Figure S2 Activating RSA neurons do not affect the anxiety level of mice**

Quantification of cumulative duration time spent in the central zone (A) and total distance mice moved (B) for the mCherry and ChR2 groups in the open-field test (n = 13 for each group)

#### Figure S3

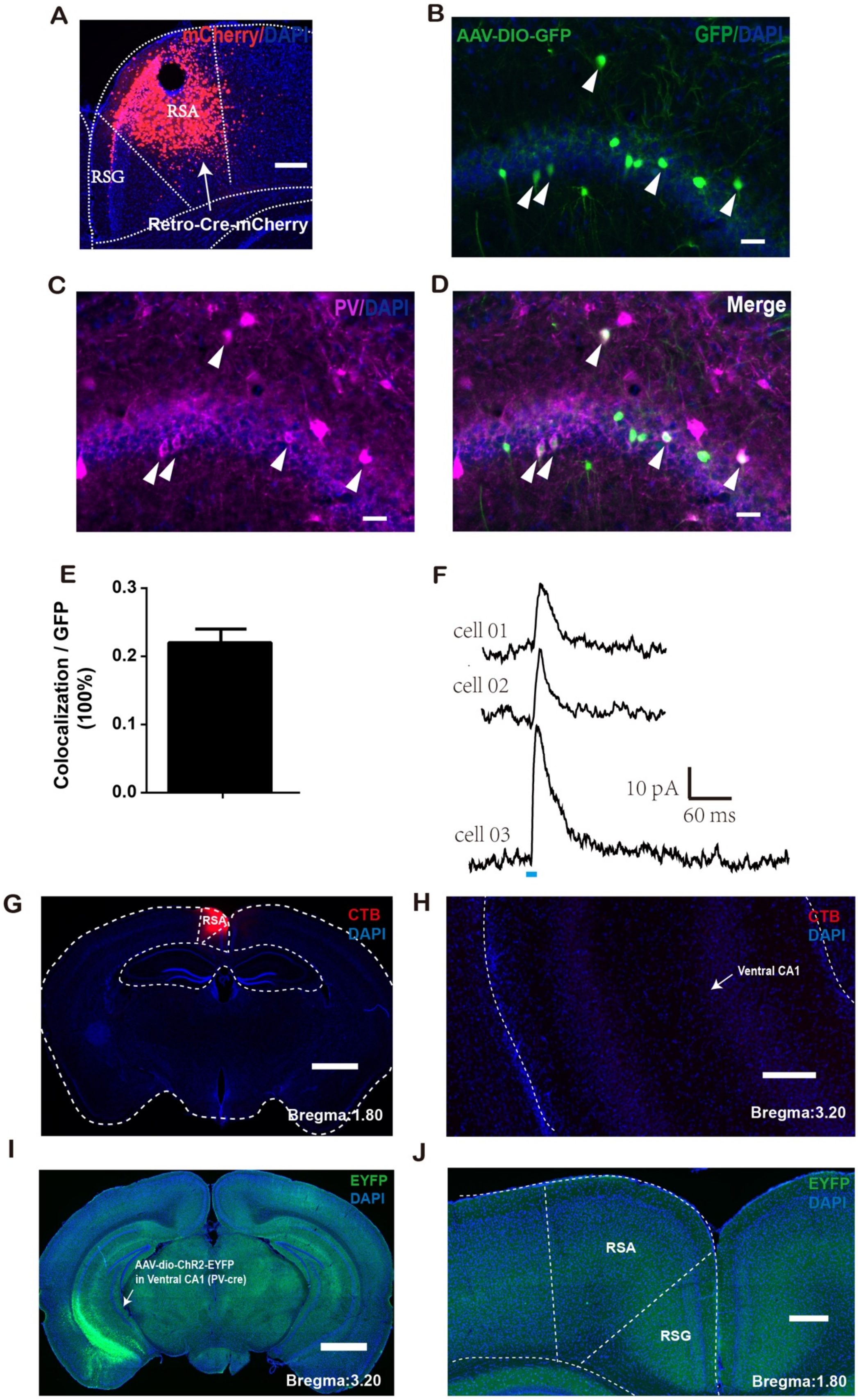


**Figure S3 Retrograde labeling of RSA-projecting CA1 PV-positive neurons**

(A) Representative images showing the inject site of AAV-Retro-Cre-mCherry in RSA; scale bar, 500μm.

The expression of GFP (B), PV (C) and merged (D) in CA1 with injecting AAV-DIO-GFP at the same time; scale bar, 20μm.

(E) Quantification of the percentage of GFP and PV double positive neurons in total GFP labeled neurons in CA1.

(G) Representative images showing the injection of CTB in RSA; scale bar, 3 mm.

(H) Representative images of ventral CA1 after CTB injection; scale bar, 1mm.

(I) Representative images showing the inject site of AAV-Dio-ChR2-EYFP in ventral CA1 of PV-cre mouse; scale bar, 3mm.

(J) Representative images of RSA after AAV-Dio-ChR2-EYFP injection; scale bar, 0.8mm.

#### Figure S4

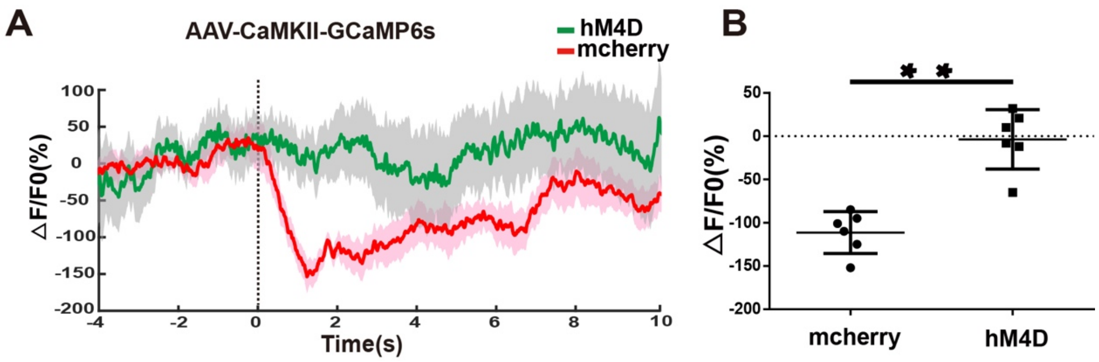


(A) Mean calcium transient associated with social interaction process. Mouse with injection AAV-CaMKII-GCaMP6s in RSA and either AAV-DIO-hM4D or AAV-DIO-mCherry in the CA1 of PV-cre mice. CNO was given through intraperitoneal injection. Solid lines indicate mean and shadow areas indicate SEM (green: hM4D, red: mCherry). Dash line at the 0s time point represents the time point when the mice actively touched the novel mice with nose.

(B) Quantification of average ΔF/F value of (0-6s) from either hM4D mice or mCherry mice (baseline is the mean value of -4s-0s) (n = 6 mice for each group).

**$p < 0.01$. Error bars represent mean ± SEM.

#### Figure S5

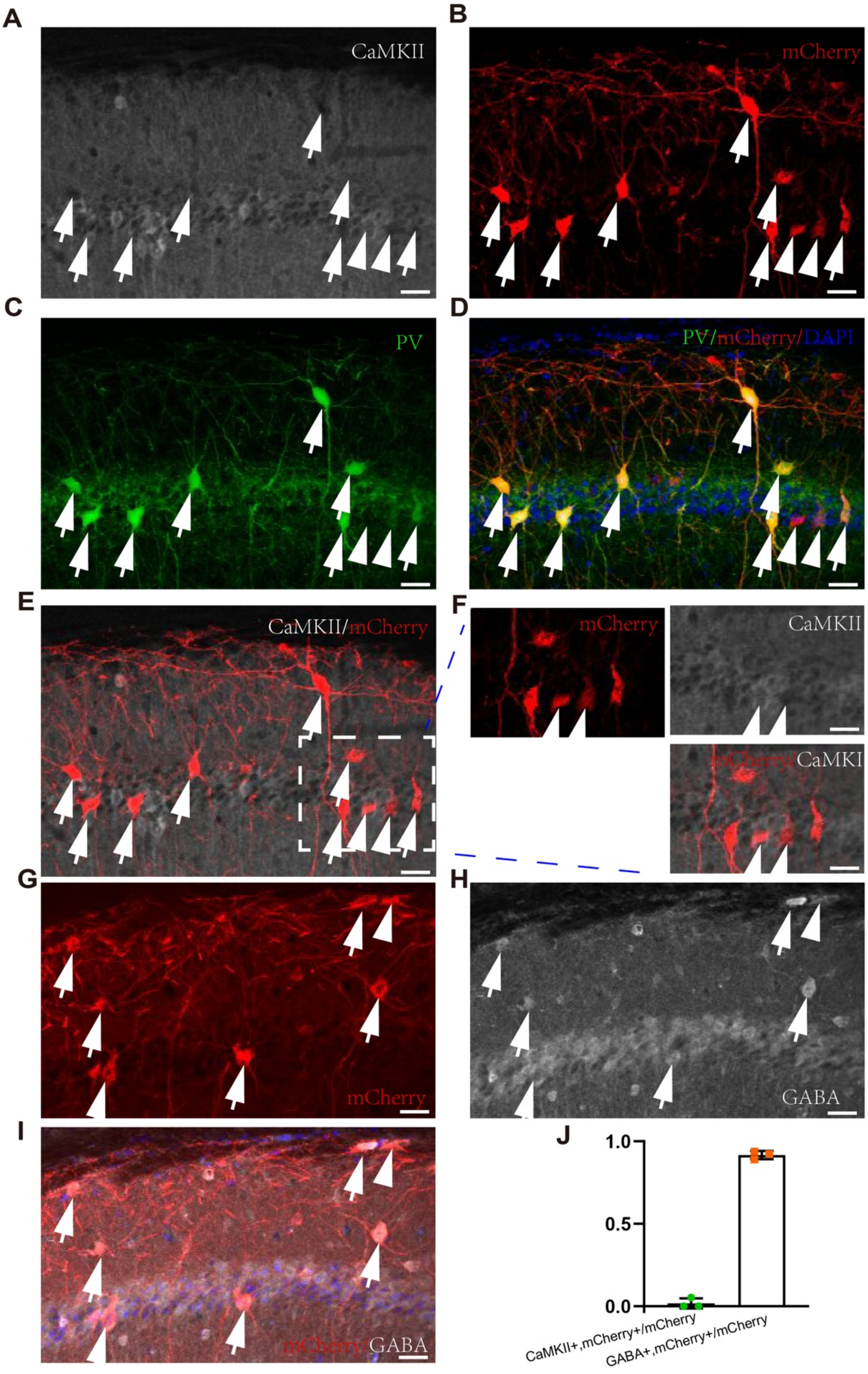


**Figure S5 Immunostaining of CA1 after AAV-pPV-mCherry injection into WT mice.**

Representative images of immunostaining in CA1 slices injected with AAV-pPV-mCherry, using antibody against CaMKII (A), mCherry (B), PV (C) respectively, and

PV + mCherry merged image (D). Arrows indicate mCherry positive neurons. Arrowheads indicate neurons with mCherry but not PV signals. Scale bar, 35μm.
(E) CaMKII + mCherry merged image. Dashed box is the area being zoom in.
(F) Zoomed in image of dashed box in (E). Scale bar, 35μm.
Representative images of immunostaining in CA1 slices injected with AAV-pPV-mCherry, using antibody against mCherry (G), GABA (H), and GABA + mCherry merged image (I). Arrow indicates mCherry positive neurons. Scale bar, 35μm.
(J) Quantification of percentage of (CaMKII, mCherry)/mCherry, and (GABA, mCherry)/mCherry neurons from (E, I).

#### Figure S6

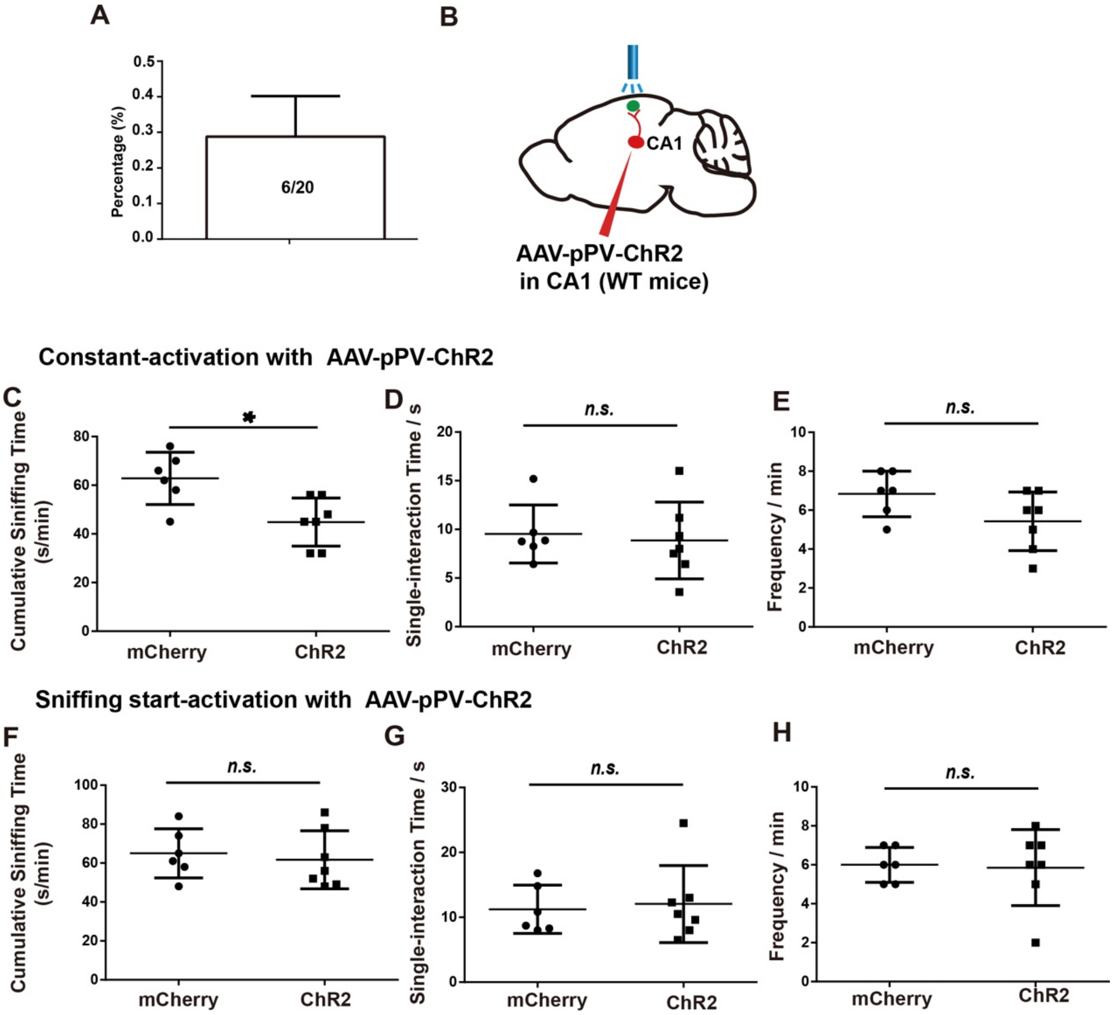


**Figure S6 AAV-pPV-ChR2 targeting CA1-PV-RSA projection in WT mice**
(A) The evoked IPSCs were recorded on 6 neurons in total 20 neurons from 3 mice.
(B) Schematic illustration for optogenetic manipulation of PV-positive input to RSA in WT mice. AAV-pPV-ChR2-mCherry was injected at CA1 of WT mice with implantation of optic stimulation fiber in RSA.

Quantification of cumulative sniffing time (C), single-interaction time (D) and sniffing frequency (E) induced by constant 473 nm light activation in the PV-ChR2 group and the PV-mCherry group.

Quantification of cumulative sniffing time (F), single-interaction time (G) and sniffing frequency (H) induced by sniffing-start 473 nm light activation in the AAV-pPV-ChR2 group and the AAV-pPV-mCherry group.

Animal number, mCherry, n = 6; ChR2, n = 7. * $p < 0.05$, Error bars represent mean ± SEM.

**Supplemental methods and materials.**

**Animals**

Mice were housed in a temperature-controlled environment (22–24°C) with ad libitum access to food and water. Mice were reared in normal lighting conditions (12-h light/dark cycle). Male and female mice (5–6 weeks old at the time of initial surgeries) from the following lines were used: C57BL/6J (The Jackson Lab，Cat# 000664), PV-Cre (B6.129P2-*Pvalb*$^{tm1(cre)Arbr}$/J, Cat #:017320), Vgat-cre (B6J.129S6(FVB)-*Slc32a1*$^{tm2(cre)Lowl}$/MwarJ, Cat#:028862), *Mef2c-het* mice were acquired from Qi Zhang's lab at ZheJiang University. The genotype of Senp1+/_ mice was determined by performing two parallel PCRs using the same forward primer from exon 8 of *Mef2c* (5'- ACTTGGCCTCTCTGCTCCACTTG-30) with different reverse primers: one primed in intron 8 of the *Mef2c* gene (50- TGTATGCTGCAAGCGTCTGTCG-30). PCR was carried out using standard techniques. All experiments were approved by the Animal Care and Use Committee of the Institute of Neuroscience, Chinese Academy of Sciences, Shanghai, China (IACUC No. NA-016-2016).

**C-fos immunostaining**

After about 12 hours' social isolation in home-cage, a stranger mouse was delivered into the cage. In the control group, a novel object was put into the cage instead. During the interaction period, the environment was kept quiet. About 1.5 hours, the mice were successively anesthetized by isoflurane, and sacrificed for C-fos immunostaining experiment.

***In vivo* optogenetic stimulation**

After about 12 hours' social isolation in home-cage, a stranger mouse was delivered into the cage. In the continuous-activation experiment, once the stranger mice were delivered into the home-cage, the optogenetic stimulation was given using blue light for activation of ChR2, with 20 Hz pulses, and the pulse width is 5ms. In the sniffing-start experiment, we turn on the light just at the beginning of the sniffing behavior and once the sniffing is over, we turn off the light immediately by hand.

**Calcium imaging**

Mice were allowed to recover for two weeks after injection of AAV virus expressing GCaMP6s. After about 12 hours' social isolation in home-cage, we put the cage with the mouse into the behavioral chamber for recording. After 5 minutes' habituation, a stranger mouse was delivered into the cage, and the calcium recording was started immediately using QAXK-FPS-TC-LED (QAXK). Onset points of the sniffing behavior were determined by analyzing the video frame-by-frame and retrieving analog signals with the MATLAB program. Photometry data were subjected to minimal processing consisting of only autofluorescence background subtraction. The values of Ca2+ transients change (ΔF/F) from-4 s to 10 s (0 s represents the onset of the actively touch of the bodies during sniffing) were derived by calculating (F-F0)/F0 for each trial, where F0 was defined as the baseline signals from -4 s to 0 s subtracted by autofluorescence background. After recording, all animals were perfused to confirm the virus expression regions and the optic fiber recording sites.

***In vivo* stereotaxic injections**

Standard stereotactic procedures were applied to mice under anesthesia (Sigma, Cat#P3761, 50mg/kg). Virus and CTB was injected with a volume of 200-400 nL/site at a rate of 20 nL/min using a micro-injector and micro-infusion pump (PHD 2000, Harvard Apparatus) to RSA according to satandard mouse brain atalas(Paxinos and Franklin Mouse Brain Atlas, 2nd edition) at the following coordinates: anteroposterior (AP), -1.82 mm; mediolateral (ML), 0.5 mm; dorsoventral (DV), -0.45 mm; to CA1 at the following coordinates: anteroposterior (AP), -1.80 mm; mediolateral (ML), 1.30 mm; dorsoventral (DV), -1.5 mm. Ten minutes after viral injection, the glass pipette was withdrew slowly to avoid the back-flow of virus. RetroAAV virus was injected with a volume of 20 nL/site at a rate of 5 nL/min. Mice were allowed 3 to 4 weeks for viral expression before the behavioral tests. The virus and titer used in this study were as follows:

AAV2/9-CaMKII-GCaMP6s-WPRE-pA (1.21E+12 v.g./mL), AAV2/9-hSyn-DIO-GCaMP6s-WPRE-pA (1.36E+12 v.g./mL), AAV2/9-CaMKII-ChR2-EYFP-WPRE-pA (1.54E+12 v.g./mL), AAV2/9-hSyn-GtACR-mCherry-WPRE-pA (1.10E+12 v.g./mL), AAV2/9-DIO-ChR2-EYFP-WPRE-pA (1.13E+12 v.g./mL), AAV2/2RetroPlus-hSyn-Cre-mCherry-WPRE-pA (1.52E+13 v.g./mL), AAV2/9-DIO-mCherry-WPRE-pA

(1.15E+12 v.g./mL), AAV2/9-DIO-hM4D-mCherry-WPRE-pA (1.35E+12 v.g./mL), AAV2/9-PV.Promoter.S5E2-hChR2(H134R)-mCherry-WPRE-pA (3.2E+11 v.g./mL), AAV2/9-PV.Promoter.E29E2-mCherry-WPRE-pA (3.1E+11 v.g./mL).

**Slice electrophysiology**

Mice were anesthetized with sodium pentobarbital (Sigma, Cat#P3761, 50mg/kg) 4 weeks after surgery. The 300 μm coronal brain slices were prepared using a vibratome (VT1200S, Leica) in an ice-cold artificial cerebrospinal fluid (aCSF) (in mM, 125 NaCl, 3KCl, 2 $CaCl_2$, 2 $MgSO_4$, 1.25 $NaH_2PO_4$, 1.3 $NaH_2PO_4$, 1.3 Na-pyruvate, 26 $NaHCO_3$, and 11 glucose, at pH = 7.4, 290-310 mOsm) saturated with 95% $O_2$ and 5% $CO_2$. After ~1hour incubation, the slice was transferred into the recording chamber which was constantly perfused with aCSF, the temperature was controlled at ~30°C by the temperature controller (Warner instrument cooperation, USA). Whole-cell recordings were performed on neurons randomly in the layer IV/V of RSA, the neurons were visualized by infrared microscope (Andor) equipped with epifluorescence and infrared-differential interference contrast (DIC) illumination. Patch pipettes were pulled from borosilicate glass (3-5 MΩ) and filled with a pipette solution consisting of (in mM), 130 K-gluconate, 20 KCl, 10 HEPES, 0.2 EGTA, 4 $Mg_2$ATP, 0.3 $Na_2$GTP, and 10 $Na_2$-phosphocreatine, at pH 7.3 (290-310 mOsm). Series resistance in whole-cell patch-clamp recording was < 30MΩ. Date were acquired with pClamp9.2 (Molecular Devices) using an AxonMultiClamp 700A amplifier (Molecular Devices), filtered at 2 kHz (low pass), and digitized at 20-100 kHz (Digidata 1322A; Molecular Devices). Before recording, the junction potential was corrected. The data analysis was performed with the Clamfit 10.3. All chemicals were purchased from Sigma. During electrophysiological recording, a pulse of 5 ms blue or yellow light from a light-emitting diode (LED) source was applied to the acute slice through an Olympus 60x water-immersion lens. The blue-light evoked responses were averaged from 6 sweeps of recording with 5s inter-sweep-interval. Recordings were made in voltage-clamp mode, with the command potential set to −70 mV to record EPSCs. It was then changed to 0 mV to record IPSCs from the same neuron. During recording in CA1, the EYFP positive neurons were identified by LED system and whole-cell recordings were performed under current-clamp mode, and a continuous blue light was given to evoke the spikes. Drugs were used in the following concentrations: SR95531-10 μM.

***In vivo* pharmacogenetic inhibition**

For inhibition of the CA1 PV axon in RSA, AAV2/9-DIO-hM4D-mCherry-WPRE-pA or AAV2/9-DIO- mCherry-WPRE-pA were unilaterally injected into the CA1, and a micro tube was implanted into the superficial layer of RSA. After 4 weeks of recovery and AAV expression, before the calcium imaging experiments, the mice were anesthetized by isoflurane, 200 nLCNO (1mM) were infused into the into the tube with a speed of 50nL/min using a micro-injector and micro-infusion pump (PHD 2000, Harvard Apparatus). 5 minutes after the infusion and the recovery from the anesthesia, the mice were used in the calcium imaging and three chamber experiment.

**Histology and immunostaining**

Mice were anaesthetized with sodium pentobarbital (Sigma, Cat#P3761, 50 mg/kg) and perfused with PBS followed by 4% PFA. After perfusion, brains were post-fixed overnight in 4% PFA at 4°C and sequentially dehydrated in 30% sucrose/PBS solution. Then brains were embedded in Optimum Cutting Temperature formulation (OCT) (SAKURA, Cat#4583) and sectioned at 40 mm with a Microtome Cryostat (Leica, CM1950) at -25°C. Floating brain sections (40 mm) were rinsed in PBS then blocked overnight at 4°C in PBS containing 5% Bovine albumin (BSA) and 0.2% Triton X-100, followed by incubating with rabbit anti-PV primary antibodies (Abcam, Cat#ab181086; 1：500) at 4°C for overnight and donkey rabbit Alexa Fluor 555 secondary antibodies (Thermo Fisher Scientific, Cat#A-31570, 1:1000) at 4°C for 2 hours. All primary and secondary antibodies were diluted with PBS containing 5% BSA and 0.4% Triton X-100. All brain sections were finally counter-stained with DAPI (Sigma, Cat#d9542, 5mg/mL, 1:1000). Sections were washed 3×10 min in PBS before incubating with secondary antibodies. For other antibody combinations, sections were rinsed with PBS, blocked and treated with primary and secondary antibodies as described above (see also KEY RESOURCES TABLE). Images were captured by objective fluorescent microscope (Olympus, VS120, 10×).

**Open field**

The mice were handled for constitutive 4 days, 4 min each time to familiarize the mice with the smell of the experimenter before the test. During the experiment, mice were put into the open field (40 × 40 $cm^2$) for 10 min. The movement of mice was recorded and analyzed by Noldus, EthoVisionXT 11.5.

**Social interaction tests**

All the mice used in the behavioral tests were male and handled for>3 days prior to behavioral tasks. The videos were recorded and analyzed by Noldus, EthoVisionXT 11.5. Before the social approach test, mice were put into the middle chamber for 10 min to habituate. In the home-cage test, a male mouse was put into a cage for about 12 hours' social isolation, a stranger mouse was delivered into the cage. During the interaction period, the environment was kept quiet. The social interaction was recorded by a camera. Then the video was analyzed manually. In the social approach test of the three-chamber test, a novel C57BL/6N male mouse (stranger) was put into the left side chambers for

10min. In the social novelty test of the three-chamber test, a novel C57BL/6N male mouse was delivered into the right chamber the video recording sustained for 10 min. The low light intensity (30Lux) was used. Then the data was recorded and analyzed by Noldus, EthoVisionXT 11.5.
